# Mechanical Stimulation via Muscle Activity is Necessary for the Maturation of Tendon Multiscale Mechanics during Embryonic Development

**DOI:** 10.1101/2021.06.29.450209

**Authors:** Benjamin E. Peterson, Rebecca A. Rolfe, Allen Kunselman, Paula Murphy, Spencer E. Szczesny

**Affiliations:** Department of Biomedical Engineering, Pennsylvania State University, United States; Department of Zoology, School of Natural Sciences, Trinity College Dublin, Ireland; Department of Public Health Science, Division of Biostatistics and Bioinformatics, Pennsylvania State University, United States; Department of Orthopaedics and Rehabilitation, Pennsylvania State University, United States

**Keywords:** Tendon development, Multiscale mechanics, mechanical stimulation, chicken embryo, collagen fibril

## Abstract

During embryonic development, tendons transform into a hypocellular tissue with robust tensile load-bearing capabilities. Previous work suggests that this mechanical transformation is due to increases in collagen fibril length and is dependent on mechanical stimulation via muscle activity. However, the relationship between changes in the microscale tissue structure and changes in macroscale tendon mechanics is still unclear. Additionally, the specific effect of mechanical stimulation on the multiscale structure-function relationships of developing tendons is also unknown. Therefore, the objective of this study was to measure the changes in tendon mechanics and structure at multiple length scales during embryonic development with and without skeletal muscle paralysis. Tensile testing of tendons from chicken embryos was performed to determine the macroscale tensile modulus as well as the magnitude of the fibril strains and interfibrillar sliding with applied tissue strain. Embryos were also treated with either decamethonium bromide or pancuronium bromide to produce rigid or flaccid paralysis. Histology was performed to assess changes in tendon size, spacing between tendon subunits, and collagen fiber diameter. We found that the increase in the macroscale modulus observed with development is accompanied by an increase in the fibril:tissue strain ratio, which is consistent with an increase in collagen fibril length. Additionally, we found that flaccid paralysis reduced the macroscale tendon modulus and the fibril:tissue strain ratio, whereas less pronounced effects that were not statistically significant were observed with rigid paralysis. Finally, skeletal paralysis also reduced the size of collagen fibril bundles (i.e., fibers). Together, these data suggest that more of the applied tissue strain is transmitted to the collagen fibrils at later embryonic ages, which leads in an increase the tendon macroscale tensile mechanics. Furthermore, our data suggest that mechanical stimulation during development is necessary to induce structural and mechanical changes at multiple physical length scales. This information provides valuable insight into the multiscale structure-function relationships of developing tendons and the importance of mechanical stimulation in producing a robust tensile load-bearing soft tissue.

## 1 Introduction

Tendons are important soft connective tissues that transfer force from muscle to bone. In general, they have strong and tough tensile load-bearing capabilities that are specialized to their anatomical location and physiological function (e.g., energy storage versus position control) (Kastelic et al., 1978; Ker et al., 2000; Thorpe et al., 2012). These unique mechanical properties arise from a complex hierarchical collagenous structure spanning multiple physical length scales over several orders-of-magnitude (Kastelic et al., 1978). While previous studies have provided insight into how this hierarchical structure (and the resulting mechanical properties) is formed (McBride et al., 1988; Birk et al., 1995; Kalson et al., 2010; Ansorge et al., 2012), several important details remain unclear. In particular, tendons undergo rapid changes in macroscale mechanics during late embryonic (in chickens) or neonatal (in mice) development (McBride et al., 1988; Ansorge et al., 2012). Previous work suggests that this mechanical transformation is due to increases in collagen fibril length (Birk et al., 1995) and mediated by mechanical stimulation via muscle activity (Pan et al., 2018). However, no study has investigated the multiscale tensile mechanics of tendons during this period of rapid change, which is necessary to fully understand the structure-function relationships of tendon development. Additionally, the biological mechanisms driving tendon maturation (including the role of mechanical stimulation) are unclear (Huang et al., 2015; Havis et al., 2016; Havis and Duprez, 2020; Tan et al., 2020). Understanding the role of mechanical stimulation in driving the rapid changes in tendon development will identify important mechanobiological principles regarding the formation of load-bearing tissues and may advance techniques for tissue engineering and repairing tendon/ligament injuries.

While the timing may vary, the overall structural and mechanical changes observed during tendon development are conserved across species. At embryonic day 10 (E10) in the chick, tendon progenitor cells organize into longitudinal columns via cadherin-mediated cell-cell junctions (Richardson et al., 2007). In this process, tenocytes organize themselves such that their cell processes form aligned longitudinal extracellular channels (Kalson et al., 2015). Short (< 50 μm) collagen fibrils are then deposited into these extracellular channels via fibripositors to form a dense and highly aligned structure (Canty et al., 2004). Up to E16, the collagen fibrils exhibit minimal increases in length despite consistent collagen deposition ((McBride et al., 1988; Birk et al., 1995). Interestingly, at E17 the collagen fibrils undergo an abrupt increase in both their length and diameter, which coincides with a rapid change in tendon macroscale mechanics that continues up to (and beyond) hatching (McBride et al., 1988). The same process is observed postnatally (P0 - P28) in mice (Ansorge et al., 2012; Kalson et al., 2015), and the coincident timing suggests that the changes in collagen fibril structure explain the changes in tendon macroscale mechanics. However, since collagen fibril lengths have not been measured in chicken embryos beyond E17, the role of this structural parameter on the mechanical properties of developing tendons is still unclear. Additionally, it is unclear how this reorganization of the collagenous network alters the local mechanical environment for tendon cells and how changes in mechanical stimulation drive tendon development.

Existing data suggest that mechanical stimulation via muscle activity plays a crucial role in driving the development and maturation of functional load-bearing tendon (Gaut and Duprez, 2016; Havis et al., 2016; Pan et al., 2018; Havis and Duprez, 2020). Mouse embryos or limb bud grafts lacking skeletal muscles show degenerated or absent tendons (Kardon, 1988), and muscle paralysis in chick embryos from E6 to E18 reduces tendon size (Germiller et al., 1998). Interestingly, motility peaks in chicken embryos starting at E12 (Wu et al., 2001) and in neonatal rats at P10 (Theodossiou et al., 2019), suggesting that muscle activity may play a particularly important role in driving the rapid structural and mechanical changes observed during late tendon development. Indeed, a recent study demonstrated that immobilizing the developing chick embryo at later stages (HH43; generally equivalent to E17) resulted in a reduction in the compressive modulus of the calcaneal tendon measured via atomic force microscopy (Pan et al., 2018). While collagen content was unchanged, muscle immobilization down-regulated the expression of the collagen crosslinker lysyl oxidase (Pan et al., 2018). However, the effect of immobilization on the length and structure of the collagen fibrils is unclear. Furthermore, mechanical testing of tendons from postnatal (P10) rats with reduced muscle activity via spinal cord transection showed an *increase* in the tensile modulus (Theodossiou et al., 2021). Therefore, it is still unclear how mechanical stimulation affects the structural maturation of collagen fibrils and the tensile mechanics of tendon during late development.

The objectives of this study were to evaluate the multiscale tensile mechanics of tendon during late embryonic chick development and to identify the effects of skeletal muscle paralysis. Specifically, we simultaneously measured the macroscale tensile mechanics and the deformations of the collagen fibrils (i.e., tensile strain and interfibrillar sliding) in tendons from chicken embryos at different developmental stages. We hypothesized that the increase in tendon macroscale mechanical properties observed during development would coincide with an increase in the fibril strains and a reduction in interfibrillar sliding, which is consistent with increasing fibril lengths (Szczesny and Elliott, 2014a). Additionally, we hypothesized that skeletal muscle paralysis would retard the changes in both the fibrillar deformations (i.e., strains and sliding) and the macroscale tensile properties observed during late embryogenesis. Finally, we hypothesized that the changes in tendon hierarchical structure (i.e., cross-sectional area, size of fibril bundles/fibers) observed during late embryonic development would also be retarded by muscle paralysis. These findings will provide valuable insight into the structural mechanisms that drive tendon development and the role of mechanical stimulation in producing a robust tensile load-bearing tissue.

## 2 Materials and Methods

### 2.1 Chick Embryo Incubation and Manipulations

Chick embryos were used for this study since their relatively large size and accessibility simplifies mechanical testing procedures and experimental perturbations of muscle activity. Fertilized eggs (White Leghorn or Ross 308) were obtained from their respective suppliers (Poultry Education and Research Center, Pennsylvania State University or Allenwood Broiler Breeders, Kildare, Ireland) and incubated at 37.7°C. All work at the Pennsylvania State University was approved by the Institutional Animal Care and Use Committee. Work on chick embryos does not require a license from the Irish Ministry of Health under European Legislation (Directive 2010/63/EU); all work was approved by the Trinity College Dublin Ethics committee.

For embryo immobilization, the eggs were windowed at E3. Briefly, 3 – 5 ml of albumin was removed from each egg using a wide gauge needle (16 ½). Scissors were then used to make a ∼2 cm diameter window in the upper surface of the egg, which was covered with transparent tape to make an airtight seal. Eggs were checked daily to remove any unviable embryos. Immobilization was induced by dripping a treatment solution on the chorioallantoic membrane through the eggshell window. Rigid paralysis of the skeletal muscles was induced by treatment with the neuromuscular blocking agent decamethonium bromide (DMB) (Sigma Aldrich) (Bowman and Rand, 1980; Mitrovic D., 1982; Osborne et al., 2002; Pitsillides, 2006). As described in **Figure 1**, three treatment regimens were used for rigid paralysis ((mild) 50 µl of 0.2% DMB on E15; (intermediate) 100 µl of 0.2% DMB at E15 followed by daily administration of 50 µl 0.2% DMB from E16 - E19; (severe) 100 µl of 0.5% DMB from E14 - E16). Flaccid paralysis was induced by treatment with pancuronium bromide (PB) (MP Biomedicals) (Reiser et al., 1988; Osborne et al., 2002; Pitsillides, 2006) via 100 µl 0.2% PB at E15 followed by daily doses of 50 µl from E16 - E19. All solutions were prepared in sterile Hank’s Balanced Salt Solution (HBSS) with 1% antibiotic/antimycotic (penicillin, streptomycin, amphotericin B; Sigma Aldrich). Control embryos were treated with HBSS and antibiotic/antimycotic alone.

**Figure 1.**
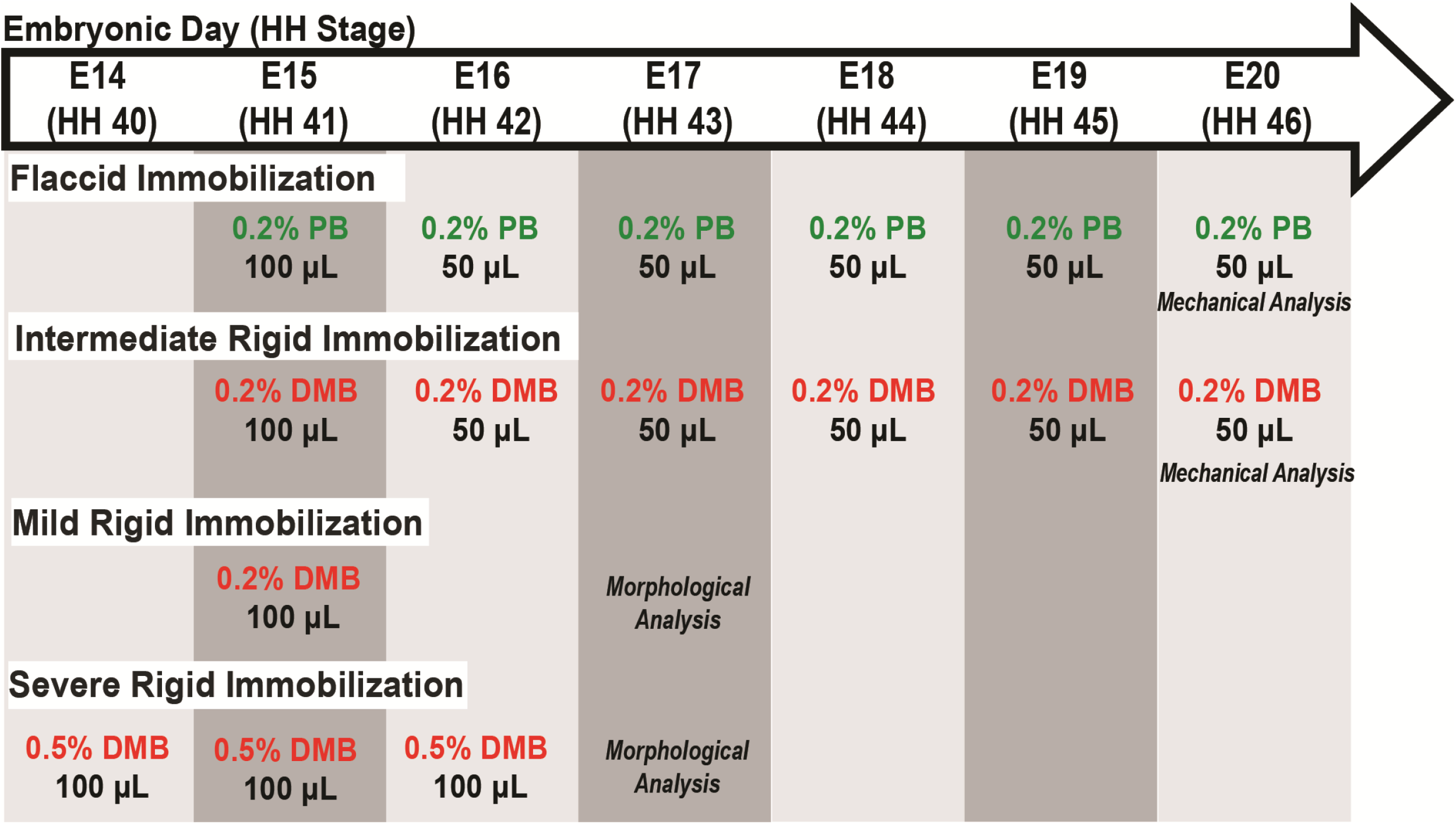
Schematic of immobilization treatment regimens, as indicated by the specific drug and their respective concentration, volumes, and embryonic day (E) in which treatment administered. Embryos sacrificed for either mechanical or morphological analysis as indicated. Abbreviations: HH; Hamburger and Hamilton stage, DMB; Decamethonium Bromide, PB; Pancuronium Bromide.

Visual inspection of embryonic motility was used to assess musculoskeletal activity during development to determine the effectiveness of the treatments (modified from Wu et al., 2001). Control, rigid paralysis, and flaccid paralysis treated embryos (n = 5 - 6) were randomly selected for motility assessments from E15 – E19. Briefly, embryos were visually examined daily through the transparent window over a 2-minute interval and any discernable individual movement (i.e., kicking or twisting) was counted. Motility counts (events / 2 mins) were averaged within each group to get a representative value for each treatment condition per day. Embryos were sacrificed either by sustained exposure to −20°C followed by cervical dislocation or by decapitation and immediate placement in ice-cold phosphate buffered saline (PBS). All embryos were staged using Hamburger and Hamilton criteria (Hamburger and Hamilton, 1951) to ensure proper developmental comparisons as represented in **Figure 1**.

### 2.2 Multiscale Mechanical Testing

#### 2.2.1 Sample Preparation

Flexor digitorum brevis (FDB) digit II tendons (**Supplemental Figure 2**) from control embryos sacrificed at E16 (HH42) (n = 6), E18 (HH44) (n = 7), and E20 (HH45/46) (n = 7) were used to assess the multiscale mechanical changes during late embryonic development. Tendons (n = 7) from intermediate DMB and PB immobilized embryos were sacrificed at E20 (HH45/46), along with vehicle controls, to investigate the effect of immobilization during the same timeframe (**Figure 1**). Hindlimbs were dissected and the FDB digit II tendon was isolated under a stereomicroscope (Nikon SMZ455T). Upon harvesting, each sample was dragged through wet lens paper with light pressure to remove the paratenon sheath and then stained with dichlorotriazinylaminofluorescein (5-DTAF, Invitrogen) by incubating in a 5 µg/ml solution of 5-DTAF and 0.1 M sodium bicarbonate buffer (pH 9.0) for 10 minutes at room temperature (Peterson and Szczesny, 2020). After incubation, samples were washed with PBS for ten minutes to remove unbound stain.

#### 2.2.2 Mechanical Testing

A 10 mm spacer was placed between two custom grips (Peterson and Szczesny, 2020) and a small amount of cyanoacrylate glue (Loctite 454) was placed on each grip face. Each stained sample was carefully lowered onto the grips such that the tarsometatarsal region of the tendon spanned the 10 mm gauge length with at least 5 mm of tissue within each grip face. Additional cyanoacrylate was added to each grip face and a drop of adhesive accelerator (Loctite 713) was used to “pot” the tissue ends. Finally, a compression plate was added to each grip and tightened to ∼ 5 inch-pounds of torque while maintaining tissue hydration with PBS.

The gripped samples were transferred to a custom uniaxial tensile-testing device mounted atop an inverted confocal microscope (Nikon A1R HD) with a PBS bath to ensure adequate hydration during testing. A preload of 0.1 g was applied, and the grip-to-grip distance was measured to establish the initial reference length. Sets of photobleached lines (PBL) (4 lines, 80 µm apart, 3 µm wide) were bleached at the sample center and ± 1.5 mm from the center (**Supplemental Figure 1**). Reference z-stack images were captured (2.5 µm z-step, 0.66 µm/px resolution) at each of the three PBL sites. The sample profile in the volumetric images was used to calculate the sample cross-sectional area (CSA) at all three PBL locations assuming an elliptical cross-section, which were averaged to determine a single CSA value. Samples were then stretched in 2% grip-to-grip strain increments at 10%/min followed by a 20-minute stress relaxation period. A prolonged relaxation period was applied to ensure that the applied stress values were stable prior to imaging (Peterson and Szczesny, 2020). At the end of each relaxation period, z-stack images were acquired at all PBL locations, and the confocal stage positions were recorded. Samples were then incrementally loaded, following the same procedure, until failure.

#### 2.2.3 Image Processing & Data Analysis

Custom image processing code (MATLAB, Mathworks) was used to determine the bulk macroscale tissue strains and the microscale fibril strains from the PBL z-stack images as previously reported (Peterson and Szczesny, 2020). Briefly, a Sobel edge detection strategy was utilized to create a two-dimensional projection of the curved tendon surface. At every position along the tissue width, the locations of the PBLs were identified as the pixel with the lowest local intensity value. For each strain increment, the microscale fibril strains were measured by tracking the displacement between the photobleached lines within each of the three PBL locations (sample center and ±1.5 mm). The fibril strains were then averaged between all three locations to generate a single representative value for each strain increment. To account for potential gripping artifacts, the macroscale bulk tissue strains were optically calculated utilizing the displacement of the peripheral (±1.5 mm) PBL sets. The fibril:tissue strain ratio was calculated at each applied strain increment by dividing the average fibril strain over the bulk tissue strain. Previous work from our lab has demonstrated that a fibril:tissue strain ratio less than one is expected for discontinuous fibrils and that the strain ratio will approach a value of one as the fibril lengths increase (Szczesny and Elliott, 2014a). The level of interfibrillar sliding was quantified by the tortuosity (i.e., waviness) of the PBLs, which was calculated as the standard deviation of the average angle of the PBLs relative to the transverse axis of the tendon. Interfibrillar sliding was calculated after each strain increment at all PBL sites and averaged to obtain a single value.

The equilibrium modulus and ultimate tensile strength (UTS) were calculated to determine the macroscale mechanical properties of the samples. A moving average (window size = 30 data points) was used to smooth the output from the load cell. The equilibrium stress value for each strain increment was calculated by averaging the last 30 seconds of data within the 20-minute stress relaxation period and was plotted versus the optically tracked bulk tissue strains. The equilibrium modulus was calculated individually for each sample by fitting a line through the second loading increment and the increment preceding the UTS in an effort to minimize the contributions of the toe and failure regions, respectively. Individual sample moduli were then averaged to get a representative modulus for the group. The UTS was defined as the peak equilibrium stress value over the course of the test.

### 2.3 Morphological Analysis

#### 2.3.1 Sectioning and Staining

Embryos were sacrificed daily from E15 (HH41) to E20 (HH46) to assess the structural changes that occur during late tendon development. Control and immobilized embryos (mild and severe treatment regimens; **Figure 1**) were sacrificed at E17 (HH43) to investigate the effects of immobilization. Briefly, the hindlimbs (distal to the knee joint) from each specimen were processed for paraffin wax sectioning. A full series of longitudinal sections (8 µm) or cross sections (8 µm) through the midpoint of the tarsometatarsal region of interest (ROI) were prepared for each specimen (**Supplemental Figure 2**). Sections were dewaxed, rehydrated, and stained to highlight connective tissue using Masson Trichrome (Sigma-Aldrich, HT15) or a fluorescently tagged natural pan-collagen binding protein (CNA35-eGFP) (Aper et al., 2014). Plasmid DNA encoding enhanced green fluorescent protein (eGFP) fused to the collagen binding protein (CNA35-eGFP) was kindly provided by the laboratory of Maarten Merkx, processed and purified as previously described (Mohammadkhah et al., 2017). Antigen retrieval was performed on histological sections using 0.01 M sodium citrate (pH 8) for 20 mins at 90°C prior to blocking of non-specific binding using 1% bovine serum albumin (BSA) in PBS for 1 h at room temperature. Slides were incubated overnight at 4°C with CNA35-eGFP (1 - 2 mg/ml) at a dilution of 1:20 - 1:50 in blocking solution. Post-antibody washes were performed in PBS and counterstained with DAPI. Histological specimens were photographed using an Olympus DP72 camera and CellSens software (v1.6). Confocal microscopy was carried out on a Leica SP8 scanning confocal using a 20X magnification objective. The confocal stack was processed and analyzed using ImageJ (v2.1.0/1.53c) software.

#### 2.3.2 Measurement of Morphological Parameters

The cross-sectional areas of the flexor digitorum longus (FDL) and the flexor digitorum brevis (FDB) digit II tendons were measured from embryos across embryonic days E15-E19 (n=2 for each, except E17, n=5) using measurements from 7-14 adjacent Masson Trichrome stained cross-sections through a 700-1300 µm portion of the medial tarsometatarsal region of interest (ROI; **Supplemental Figure 2**). The effects of immobilization were assessed at E17 (n=3 for control, severe and mild treatment regimens), taking measurements from 5-11 adjacent sections per specimen. The boundary of collagen positive (blue) staining at the bundle edge was defined as the boundary of the tendon for quantification. The cross-sectional area of the FDL and FDB tendons were quantified independently using ImageJ (v2.1.0/1.53c).

The same medial ROI was used to assess the spacing between the FDL and FDB tendons over time and following immobilization (**Supplementary Figure 2**). For each section from each sample, 6-10 measurements were taken across the extent of the interface (where approximately parallel) between the tendons, and averaged, to represent the spacing between the tendons (**Supplemental Figure 2C**; red lines indicate typical regions for measurement).

**Figure 2.**
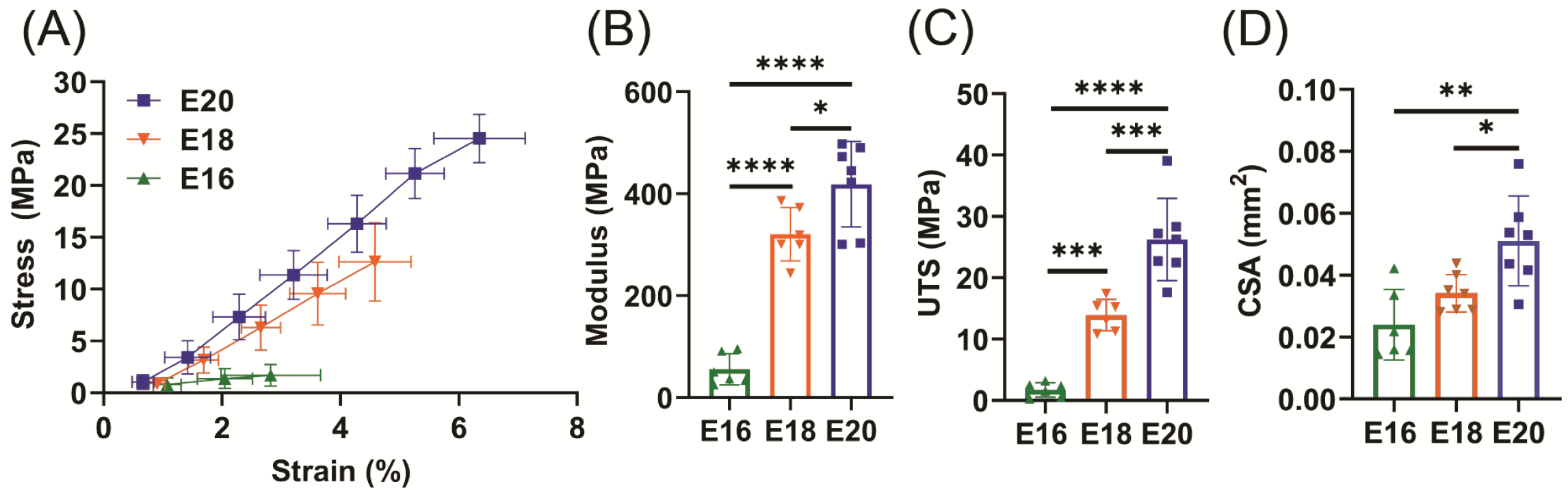
Flexor digitorum brevis (FDB) digit II tendon macroscale mechanical response with maturation. (A) Equilibrium stress vs strain response for E16 (n = 6), E18 (n =7), and E20 (n = 7) timepoints. (B) Significant increase in equilibrium modulus response with embryonic maturation. (C) Significant increase in ultimate tensile stress (UTS) with development. (D) A significant increase in the cross sectional (CSA) of the tendon was observed with maturation when measured under confocal microscopy. * p < 0.05, ** p < 0.01, *** p < 0.001, **** p < 0.0001.

The collagen fiber diameters were estimated from confocal fluorescence volumetric images of the CNA35-eGFP stain across developmental stages and following immobilization. Two to three image planes from 5-9 ROIs (100 µm^2^) per specimen were analyzed, which provided a measurement of 79-117 individual fibers per specimen (immobilization experiment (E17): n= 6 control, n= 4 severely immobilized, n= 2 mildly immobilized) (**Figure 1**). Individual fibers in the 100 µm^2^ ROI were measured using ImageJ software from edge to edge (**Supplemental Figure 2**) as determined by a higher intensity of CNA35-eGFP fluorescence.

### 2.3 Statistical Analysis

Significance was set at p ≤ 0.05 for all tests. For mechanical characterization, statistical analysis was conducted using SAS (9.4) and GraphPad Prism (8.3.0). A one-way ANOVA was conducted to evaluate the UTS and equilibrium modulus with respect to development or paralysis treatment. Post-hoc Tukey’s tests corrected for multiple comparisons were conducted to evaluate the UTS and modulus with increased maturation. A post-hoc Dunnett’s test corrected for multiple comparisons evaluated the effect of immobilization treatments compared to the vehicle control. An ANCOVA with Bonferroni post-hoc tests was used to determine if the embryonic motility was affected by development or immobilization treatment. Differences in the fibril:tissue strain ratio and interfibrillar sliding were evaluated using a general linear model with correlated errors that takes into consideration the correlated strain increments per specimen as a covariate. A linear-regression was conducted to test whether the fibril:tissue strain ratio was negatively correlated with applied tissue strain and a one-sample t-test was conducted to determine if the group mean was significantly less than 1. Similarly, a linear regression was conducted to test whether the fibril sliding behavior was positively correlated with applied tissue strain. For morphological analysis, differences in tendon area, distances between tendons, and fiber diameter were assessed by univariate ANOVA followed by Tukey post-hoc tests using SPSS (SPSS Statistics v26, IBM).

## 3. Results

### 3.1 Changes in Tendon Multiscale Mechanics and Structure with Development

There was a significant increase in the macroscale modulus with development (**Figure 2B**) (p < 0.0001) with significant differences observed between E16 and both E18 (p < 0.0001) and E20 (p < 0.0001) as well as between the E18 and E20 timepoints (p < 0.05). The ultimate tensile stress (UTS) was also significantly increased at later developmental timepoints (Figure **2C**) (p < 0.0001) with similar individual differences between the E16 and both E18 (p < 0.001) and E20 timepoints (p < 0.0001) as well as between the E18 and E20 timepoints (p < 0.001). Finally, there was a significant increase in the tendon cross-sectional area as measured by confocal microscopy with development (**Figure 2D**) (p < 0.01). Post-hoc comparisons found a significant difference between E16 and E20 (p < 0.01) as well as between E18 and E20 (p < 0.05) but no difference the E16 and E18 samples. These large changes in macroscale mechanics are consistent with prior data showing increases in mechanical properties during late tendon development (McBride et al., 1988).

To assess the mechanisms underlying these macroscale mechanical changes during development, we measured the fibril:tissue strain ratio and interfibrillar sliding as a function of applied tissue strain. Previous work has demonstrated that a fibril:tissue strain ratio less than one is expected for discontinuous fibrils and that the fibril:tissue strain ratio will increase (while the interfibrillar sliding will decrease) as the collagen fibrils elongate (Szczesny and Elliott, 2014b). All embryonic samples, regardless of their developmental timepoint, displayed a mean fibril:tissue strain ratio that was less than one and negatively correlated with the applied tissue strain (**Figure 3A**) (Control: p < 0.05, PB: p < 0.0001, DMB, p < 0.001). A general linear model found a significant increase in the fibril:tissue strain ratio between E16 and both the E18 (p < 0.05) and E20 (p < 0.001) timepoints. Additionally, the interfibrillar sliding was positively correlated with the applied tissue strains for all samples (**Figure 3B**). However, there were no significant difference in the interfibrillar sliding with development.

**Figure 3.**
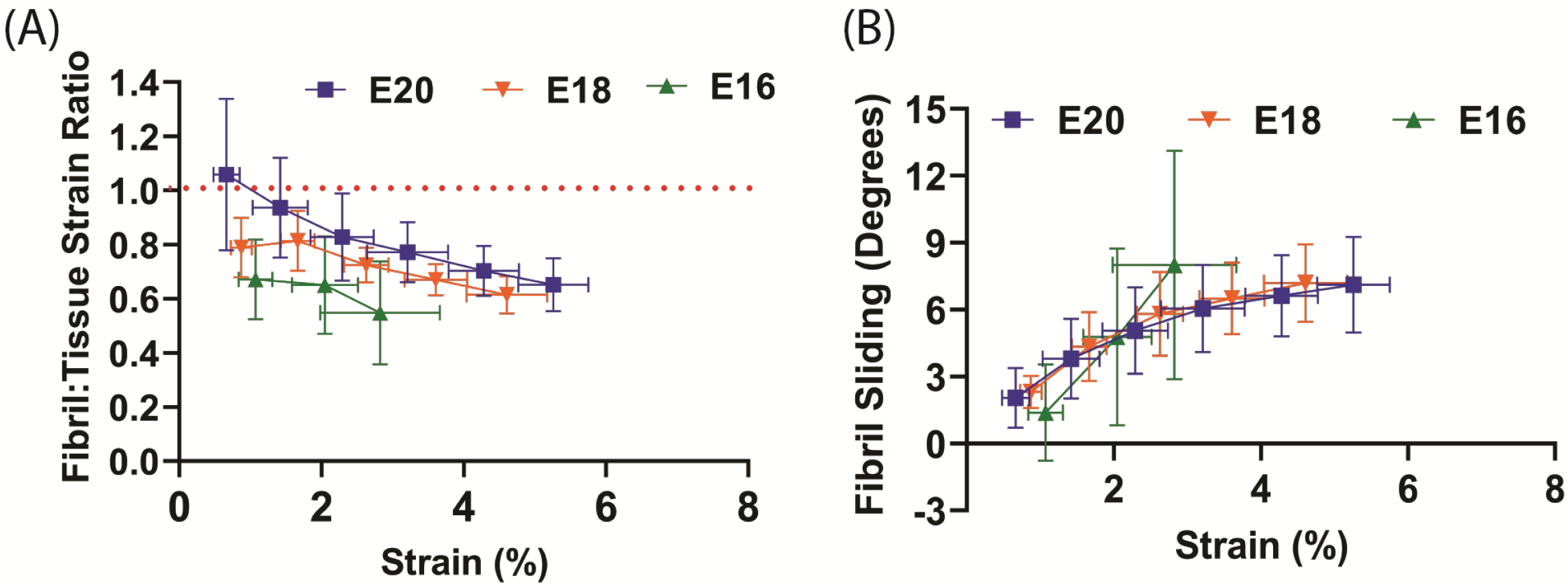
Multiscale mechanical response of the flexor digitorum brevis (FDB) digit II tendon as a function of applied tissue strain with increasing development. (A) A significant increase in the fibril: tissue strain ratio was observed between E16 vs. E18 (p < 0.05) and E16 vs. E20 (p < 0.001). (B) No significant difference in the fibril sliding behavior was observed with developmental age.

To profile the morphological changes in the FDL and FDB tendons over the developmental time window, transverse histological sections through the tendon midpoint were analyzed (**Supplementary data Figure 2; Figure 4A**). An increase in tendon size was observed through measurements of the cross-sectional area (**Figure 4B**); in both tendons there was a steady increase in the mean value during development with a larger step increase at E17, particularly for the FDL. The distance (spacing) between the tendons did not change significantly over time, but there was a visible increase between E15 and E16 (**Figure 4A**), reflected in the mean distance measurement (**Figure 4C**). Longitudinal sections, stained with the fluorescently tagged pan-collagen binding protein CNA35-eGFP, allowed visualization of the underlying collagen network across time **(Figure 4D**). The fibril bundles (fibers) visibly enlarged, reflected in an incremental increase in the average fibril bundle diameter over time (**Figure 5E**). There was no obvious change in the alignment of the fibril bundles over time, at least at this level of resolution.

**Figure 4.**
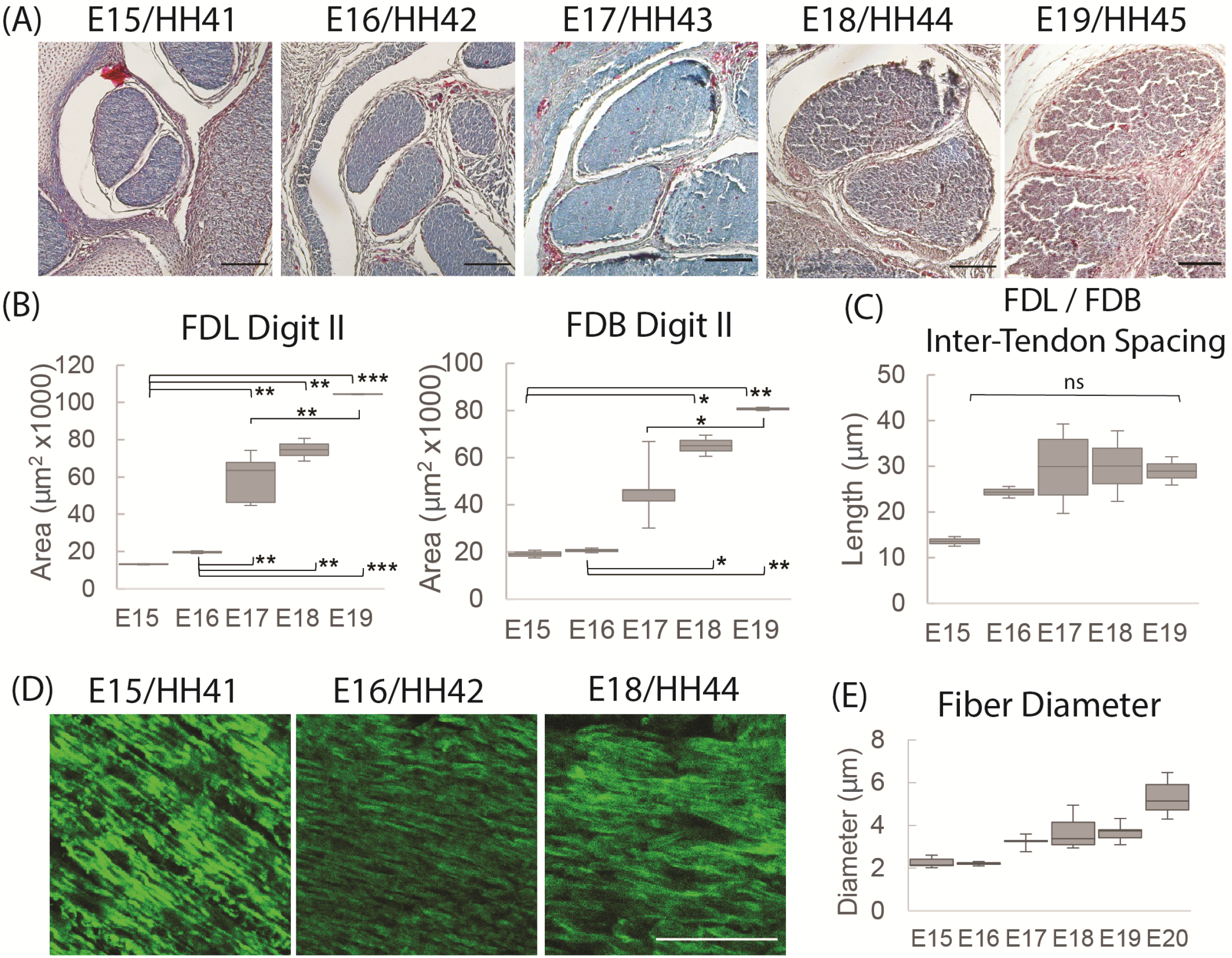
Structural analysis of tendons between E15 (HH41) and E19 (HH45). (A) Cross-sectional profile of the flexor digitorum longus (FDL) and brevis (FDB) digit II tendons stained with Masson Trichrome in the tarsometatarsal region of the developing chick hindlimb over embryonic days, as indicated. Scale bar 100μm. (B) Measurements of cross-sectional area of the FDL and FDB digit II tendon over developmental time. (C) Measurements of spacing between the FDL and FDB digit II tendons over developmental time. (D) Images of longitudinal tendon fibers in the tarsometatarsal region stained with collagen binding protein over developmental time. Scale bar 50μm. (E) Measurement of tendon fiber diameter over time. For (B) and (C): n=2 per day except E17 n=5; For (E) n=1 (multiple sections from 1 specimen/stage measured). * p ≤ 0.05, ** p ≤ 0.01, *** p ≤ 0.001.

**Figure 5.**
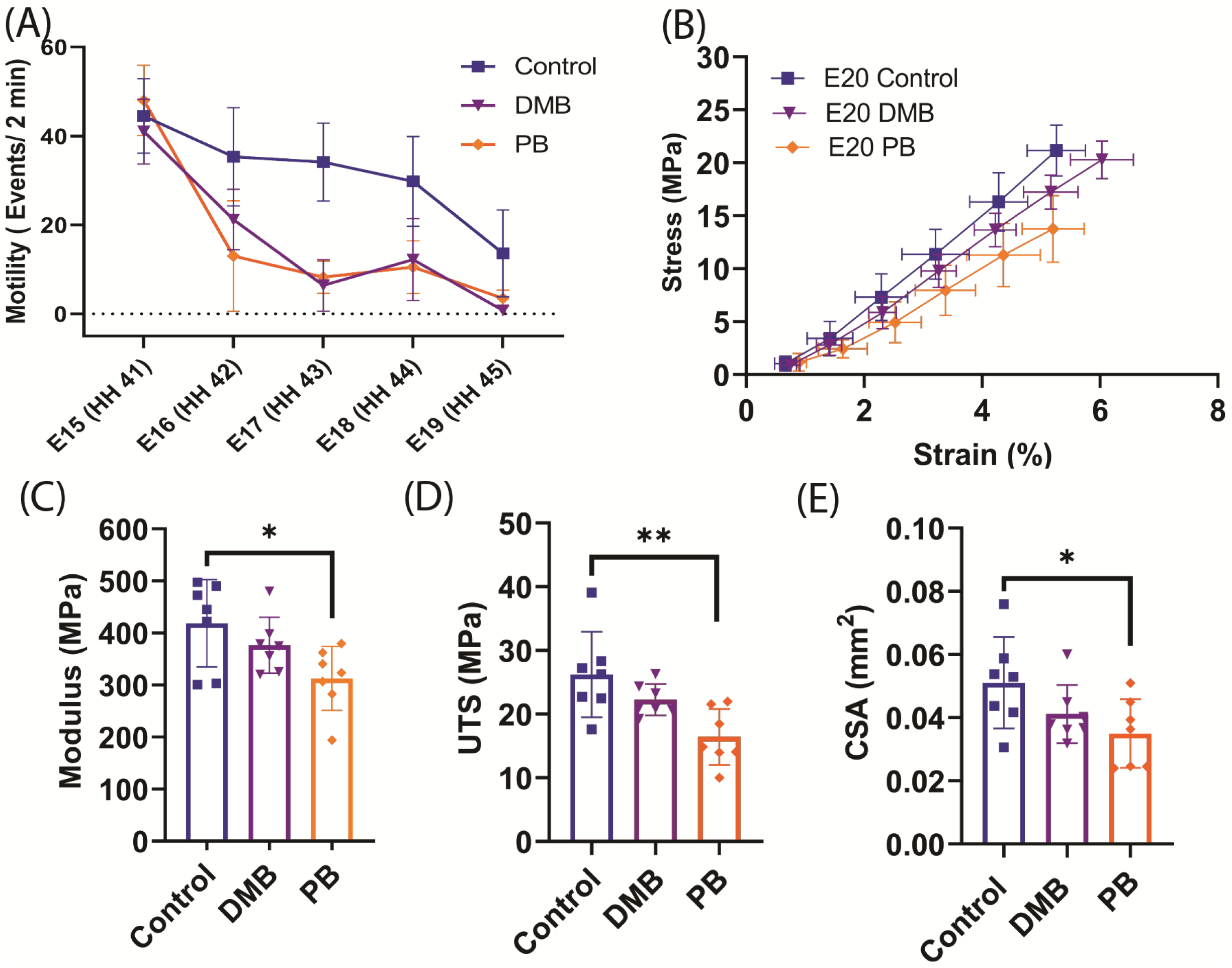
Muscle paralysis and its effect on the macroscale mechanical properties of the flexor digitorum brevis (FDB) digit II tendon during development. (A) Paralysis treatments and saline control were conducted from E15 (HH41) - E19 (HH45). Significant decrease in motility observed under rigid (DMB, p < 0.05) and flaccid (PB, p < 0.05) paralysis. No significant difference in motility counts between the two paralysis methods. (B) Equilibrium stress vs. applied strain for E20 (HH46) samples after control (n=7), rigid paralysis (DMB, n=7), and flaccid paralysis (PB, n=7) treatments. (C) Equilibrium modulus is significantly reduced with flaccid paralysis treatment (p < 0.05). (D) Significant decrease in ultimate tensile stress (UTS) with flaccid paralysis (p < 0.01). (E) Cross-sectional area of flexor digitorum brevis (FDB) tendon was observed under flaccid paralysis conditions when measured under confocal microscope. (p < 0.05). * p ≤ 0.05, ** p ≤ 0.01

### 3.2 Effect of Muscle Paralysis on Tendon Development

Rigid and flaccid paralysis treatment, with DMB (intermediate regimen) and PB, respectively, significantly reduced embryonic motility through the target period (DMB: p < 0.05, PB p < 0.05) (**Figure 5A**). No significant difference in motility was observed between the rigid DMB or flaccid PB paralysis treatments (p = 0.67). After validating the immobilization treatments, we investigated their effect on the mechanical and structural changes that occur during late embryonic development (**Figure 2 & 3**). At the macroscale level, flaccid paralysis (PB treatment) significantly reduced the equilibrium modulus compared to vehicle control samples (**Figure 5C**) (p < 0.05). Similarly, flaccid paralysis significantly decreased both the UTS (p < 0.01) and CSA (p < 0.05) (**Figure 5D&E**). No significant differences in the macroscale mechanical properties were observed under the rigid paralysis (DMB) conditions compared to controls (modulus: p = 0.47, UTS: p = 0.24, CSA: p= 0.22).

To investigate how the multiscale mechanics were perturbed under paralysis conditions, we measured the fibril:tissue strain ratio and interfibrillar sliding as a function of applied tissue strain for each treatment group. All samples (control, PB, and DMB) displayed a mean fibril:tissue strain ratio that was less than one (p < 0.05) and was negatively correlated with applied tissue strain (**Figure 6A**) (p < 0.001). A general linear model found a significant decrease in the fibril:tissue strain ratio for the flaccid (PB) immobilized samples compared to control tissues (p < 0.05). No significant differences were found between the rigid immobilized and control tissue (p = 0.11). The interfibrillar sliding was positively correlated with the applied tissue strain for all samples (**Figure 6B**). However, no significant group effects were observed under paralysis conditions in comparison to the control (PB: p = 0.10, DMB: p = 0.78).

**Figure 6.**
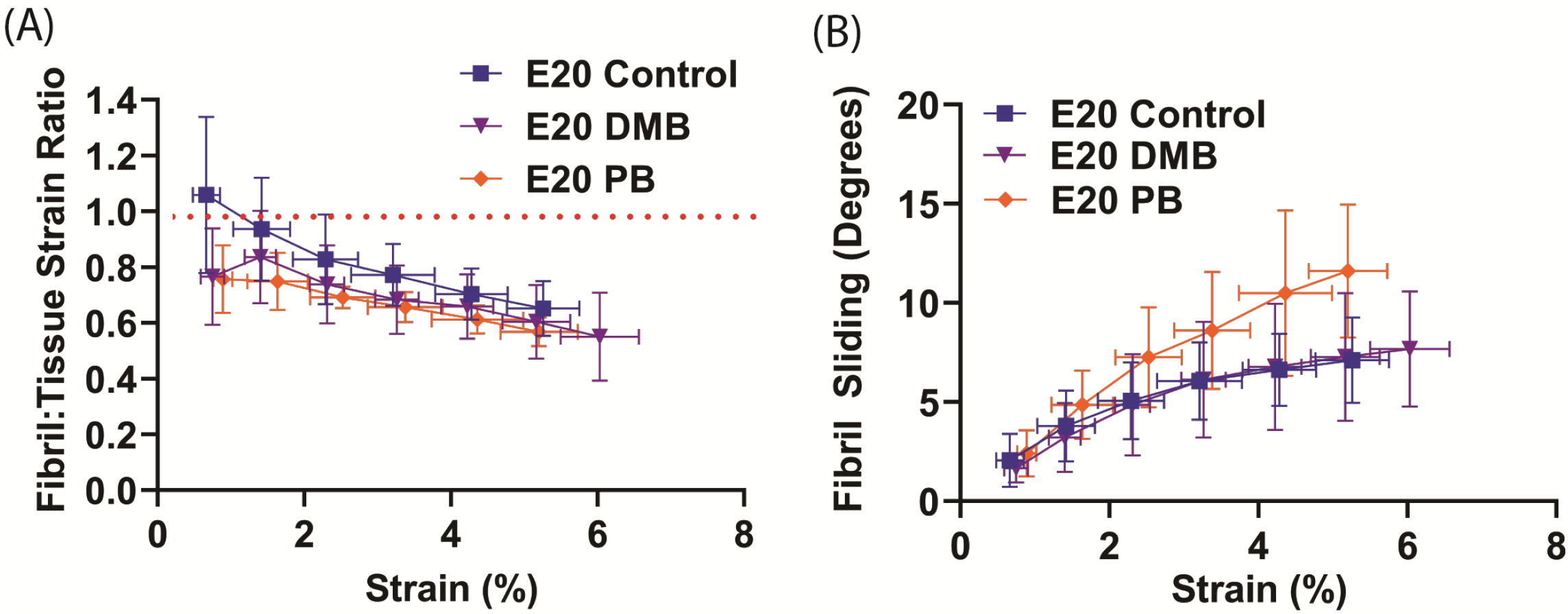
Multiscale mechanical response at E20 as a function of applied tissue strain after immobilization treatments. (A) The fibril:tissue strain ratio was significantly lower with flaccid paralysis (PB) treatments in comparison to the control tissue (p < 0.05). No significant difference was observed between the rigid paralysis (DMB) and the vehicle control tissue (p = 0.106). (B) All samples displayed an increase in fibril sliding angle with applied stress. No significant group effects were observed under paralysis conditions in comparison to vehicle controls (PB vs. Control: p = 0.10, DMB vs. Control: p = 0.78).

Rigid paralysis of chick embryos caused multiple changes to the morphology of the FDL and FDB tendons at E17 (**Figure 7**). Two regimens of DMB treatment were tested for effect; a more severe treatment that involved 0.5% applied daily from E14 and a milder regimen consisting of a single treatment with 0.2% at E15 (**Figure 1**). At a gross morphological level, both the cross-sectional area and the spacing between tendons were significantly reduced with the more severe exposure alone (p < 0.01). However, a greater sensitivity to treatment was observed at a smaller length scale, with fiber diameter significantly reduced by both mild and severe treatments.

**Figure 7.**
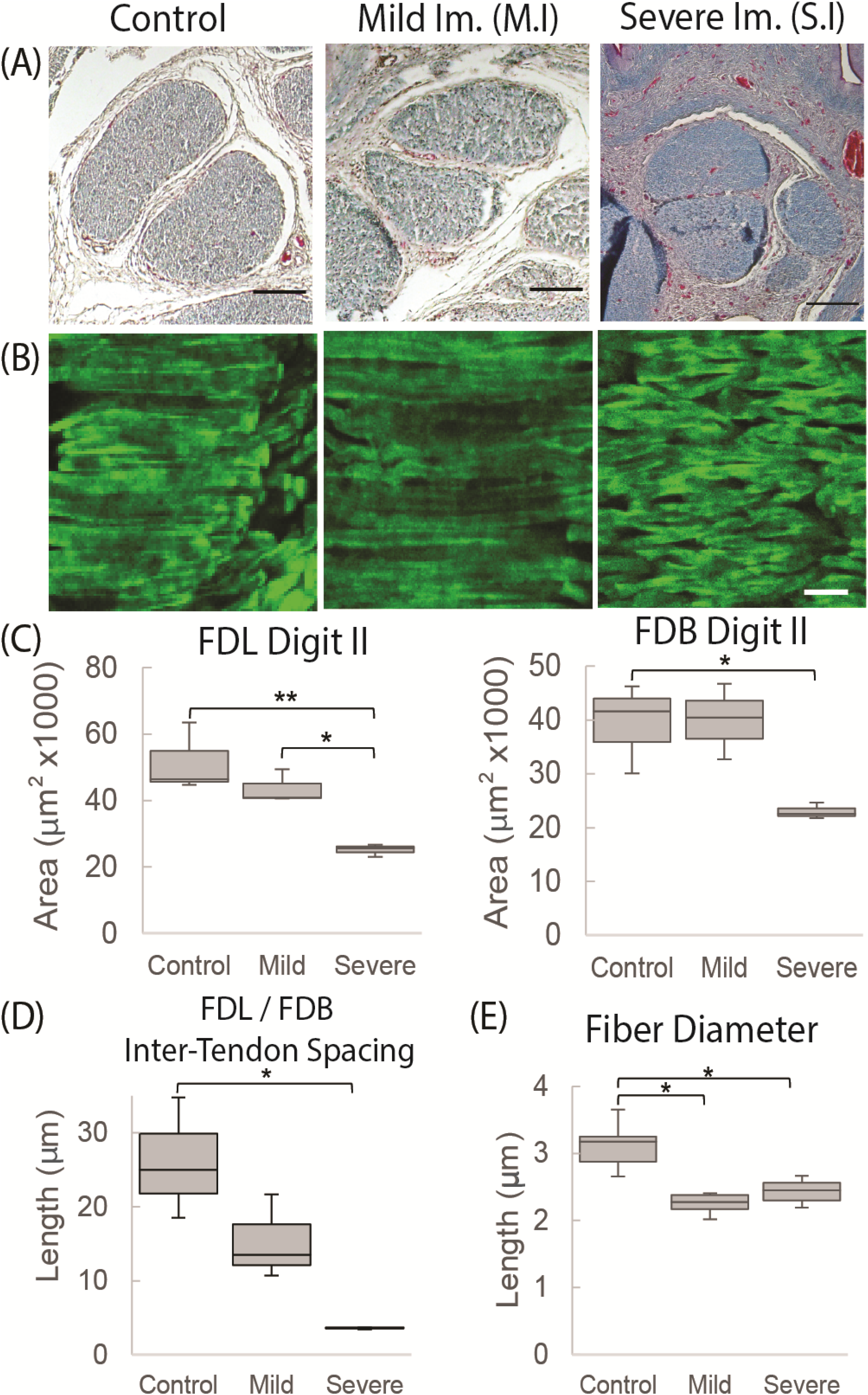
Structural organization of tendons at E17 is altered due to rigid paralysis. (A) Representative cross sections through the flexor digitorium longus (FDL) and brevis (FDB) digit II tendons stained with Masson Trichrome in controls and following mild (M.I) and severe immobilization (S.I) DMB treatments (Figure 1). Scale bar 100μm. (B) Representative images of longitudinal sections stained with collagen binding protein (CNA35-eGFP). Scale bar 10μm. (C) Cross-sectional area of both tendons is reduced significantly following the more severe immobilization regimen. (D) Spacing between tendons is significantly reduced following the more severe immobilization regimen. (E) Fiber diameter within the tendons is significantly reduced following both mild and severe immobilization regimens. n=3, all groups in C and D; n=6 (control), n=4 (S.I), n=2 (M.I) in E. *p≤0.05, **p ≤0.01

## 4. Discussion

This study demonstrated that there is a pivotal shift within tendon structure-function relationships during late stages of embryonic development that facilitates the functional load-bearing capabilities of tendons. Specifically, there is an increase in the fibril:tissue strain ratio after E16 in chicken embryos (**Figure 3A**), which is consistent with an increase in the length of the collagen fibrils (Birk et al., 1995; Szczesny and Elliott, 2014b) and the increase in macroscale tensile mechanics (**Figure 2**) (McBride et al., 1988). It is also important to note that the E20 tissue still displayed a negatively correlated fibril:tissue strain ratio at larger applied strains, which is consistent with discontinuous fibrils and not progressive breakage of continuous fibrils (Peterson and Szczesny, 2020). However, the observed upward shift in the fibril:tissue strain ratio with maturation indicates a transition in the collagenous loading behavior, in which strains are being transferred to the collagen fibrils more directly. While shorter fibrils should exhibit more relative sliding with tissue stretch (Szczesny and Elliott, 2014a, 2014b), no significant difference in interfibrillar sliding was observed between E16 and the later developmental timepoints. Nevertheless, there was an increase in the standard deviation within the E16 group, which is possibly due to variable interfibrillar load-transfer capabilities that is representative of a developmental stage undergoing rapid ultrastructural changes (Birk et al., 1995). This is further supported by morphological analysis of the FDB and FDL tendons, which showed a significant increase in the CSA (**Figure 4B**) and an incremental increase in the average fiber diameter with development in the samples assayed (**Figure 4E**). Together, this study supports our hypothesis that there is a reorganization of the collagenous network during late stages of tenogenesis that increases the strain transmitted to the collagen fibrils and generates a rapid increase in tendon mechanical properties.

Our findings also demonstrate that the observed changes in fibril loading and macroscale tensile mechanics are impeded by musculoskeletal paralysis, suggesting that mechanical stimulation during late stages of tenogenesis is critical to mediating developmental changes in the collagen fibril network. Specifically, flaccid muscle paralysis reduced the macroscale modulus, UTS, and CSA (**Figure 5**) as well as the fibril:tissue strain ratio (**Figure 6**) with a trending increase in the interfibrillar sliding (p = 0.1). Interestingly, the effect of rigid paralysis on the macroscale mechanics (**Figure 5B - E**) and fibril loading (**Figure 6A**) was not significant, despite a comparable decrease in embryo movement as flaccid paralysis treatment (**Figure 5A**). This variation in response is likely attributed to the different modes of action in which these drugs induce immobilization. PB competitively blocks the acetylcholine receptor at the motor endplate to inhibit muscle contraction, causing flaccid or limp paralysis (Bowman and Rand, 1980; Osborne et al., 2002). In contrast, DMB irreversibly binds to acetylcholine receptors in the motor endplate to induce permanent contraction of muscle, generating a prolonged static load across the tissue (Bowman and Rand, 1980; Osborne et al., 2002). We hypothesize that while both immobilization models reduce cyclical musculoskeletal stimulation, the static loading environment induced by the DMB treatment reduces the overall phenotypic effect. Differences between the effects of rigid and flaccid paralysis on skeletal development have also been noted. Conversely, the effects of rigid paralysis on the spine were more pronounced (Rolfe et al., 2017), as was the effect on length and breadth of the cartilaginous skeletal rudiments, while flaccid paralysis caused greater disruption of post-cavitation stage joints (Osborne et al., 2002). Although the static loading experienced by tissues under rigid paralysis (Nowlan et al., 2008) can have a more detrimental effect under some circumstances, it is clearly not the case for these stages of tendon development. Despite this, rigid paralysis treatments still have a significant effect on the tissue morphology in a dose dependent manner (**Figure 7**). Histological characterization demonstrated that severe rigid immobilization was required to induce a significant reduction in the CSA and inter-tendon spacing, with minimal effects under a mild treatment paradigm. Interestingly, both the mild and severe rigid immobilization treatments induced comparable reductions in the collagen fiber diameter, suggesting that ultrastructural changes in the collagen network are highly sensitive to the loss of external mechanical stimulation, which is consistent with our mechanical data using a similarly mild treatment (**Figures 5 & 6**).

While our data suggest that mechanical stimulation plays an important role in driving the structural and mechanical changes observed during late tendon development, the biological mechanisms mediating this are unclear. Prior work has shown that the loss of mechanical stimulation during late stages of tendon development reduces LOX expression, inhibiting cross-linking activity within the collagenous network and reducing the tissue compressive modulus (Makris et al., 2014; Pan et al., 2018). However, numerous other ECM components are known to regulate collagen fibril interconnections, and it is not clear how they are affected by mechanical stimulation. For example, decorin and biglycan bind to the surface of collagen fibrils and inhibit collagen fibril fusion (Schönherr et al., 1995; Graham et al., 2000; Zhang et al., 2006), which may explain the rapid change in collagen fibril structure during late tendon development (Robinson et al., 2005; Zhang et al., 2006). Furthermore, their relative expression decreases during development precisely when fibril lengths/diameters grow rapidly (Birk et al., 1995, 1995; Chen and Birk, 2013). However, previous work indicates that decorin expression is positively correlated to mechanical stimulation (Hayashi et al., 2019). This suggests that the reduction in the fibril:tissue strain ratio (potentially due to a retardation in collagen fibril elongation) seen with flaccid paralysis (**Figure 7A**) may not be due to a change in decorin or biglycan expression. Another important note is that there were still significant increases in tendon macroscale mechanics with PB treatment when compared to normally developing E16 tissue, suggesting that some load-bearing elements may develop independently of mechanosensitive mechanisms. Further investigations are required to characterize the changes in tendon structure-function relationships during late tendon development, its sensitivity to musculoskeletal stimulation, and other regulatory mechanisms that link these two. Overall, this study supports our hypothesis that mechanical stimulation during late tendon development plays a critical role in mediating the gross structural and mechanical changes across multiple length scales.

A limitation to this study is that it took 48 hours to produce the full immobilization effects of the paralytic drug treatments. Therefore, it is possible that some mechanosensitive maturation occurred during E15 - E17 even with treatment, which could explain the increase in mechanical properties of the E20 PB treated samples compared to normally developing E16 tissue. An additional limitation is that confocal microscopy does not have the optical resolution required to directly visualize collagen fibrils. That is, our imaging resolution is 0.66 um/px, while the average diameter of collagen fibrils during these developmental timepoints range from 45 nm to 60 nm (McBride et al., 1988). However, fibril bundles (i.e., fibers) during this same developmental window are approximately 2 - 5 µm in diameter (Silver et al., 2003), suggesting that we are capturing a localized average fibril response. Finally, changes in fibril structure (e.g., fibril length, diameter, and interfibrillar spacing) and their dependence on musculoskeletal activity during late tendon development were not directly measured. Nevertheless, this study provides valuable insight into the multiscale structure-function relationships of developing tendons and the importance of mechanical stimulation in producing a robust tensile load-bearing soft tissue.

## 5. Acknowledgments

We thank Dr. Maarten Merkx and his lab for providing the plasmid DNA encoding eGFP fused to CNA35 used to generate the collagen binding probe.

## 6. Funding

This work was supported by the National Institutes of Health [R21 AR075941].

## 7. Author Contributions

Benjamin Peterson and Rebecca Rolfe carried out the reported experiments and data analysis. Allen Kunselman aided in the design of the statistical analysis. Spencer Szczesny and Paula Murphy supervised the work and contributed to the interpretations of the results and writing of the manuscript. Benjamin Peterson was the primary author of the manuscript. All authors provided crucial feedback to shape the research, analysis, and manuscript.

## 8. Conflicts of Interest

The authors declare no competing financial interests.

## Contribution to the field

Tendons are connective tissues that facilitate the transmission of force between muscle and bone. The high load-bearing capabilities of tendons are established during late-stages of development (chick) and are believed to be mediated by structural transformations within the hierarchical organization of collagen. Prior work has suggested that this structure-function transformation depends on mechanical stimulation during development. However, no study has investigated the multiscale structure-function relationships during this critical developmental period or established how this transformation is sensitive to mechanical stimulation. Therefore, the objective of this study was to evaluate changes in tendon mechanics and structure at multiple length scales during embryonic development with and without skeletal muscle paralysis. Our study identified a pivotal shift in the loading behavior during late-stages of tenogenesis, characterized by an increase in strain transmission to collagen fibrils that generates a rapid increase in tendon mechanical properties. We further identified that this structure-function transformation is impeded under musculoskeletal paralysis, indicating that mechanobiological activity plays a crucial role in mediating the functional capabilities of tendons. Understanding the dynamic loading conditions during development is essential to identifying mechanobiological processes that dictate the formation of tendons and may advance tissue engineering and regenerative medicine approaches for tendon/ligament injuries.

## Ethics statements

### Studies involving animal subjects

#### Generated Statement

The animal study was reviewed and approved by Pennsylvania State University Institutional Animal Care and Use Committee and the

Trinity College Dublin Ethics Committee.

### Studies involving human subjects

#### Generated Statement

No human studies are presented in this manuscript.

### Inclusion of identifiable human data

#### Generated Statement

No potentially identifiable human images or data is presented in this study.

### Data availability statement

#### Generated Statement

The raw data supporting the conclusions of this article will be made available by the authors, without undue reservation.

## Notes

*Conflict of interest statement* The authors declare that the research was conducted in the absence of any commercial or financial relationships that could be construed as a potential conflict of interest

### Competing Interest Statement

The authors have declared no competing interest.

