## Supplemental Figures 1&2 for "Mechanical Stimulation via Muscle Activity is Necessary for the Maturation of Tendon Multiscale Mechanics during Embryonic Development"

### *Supplementary Material*

#### 1. Supplementary Figures

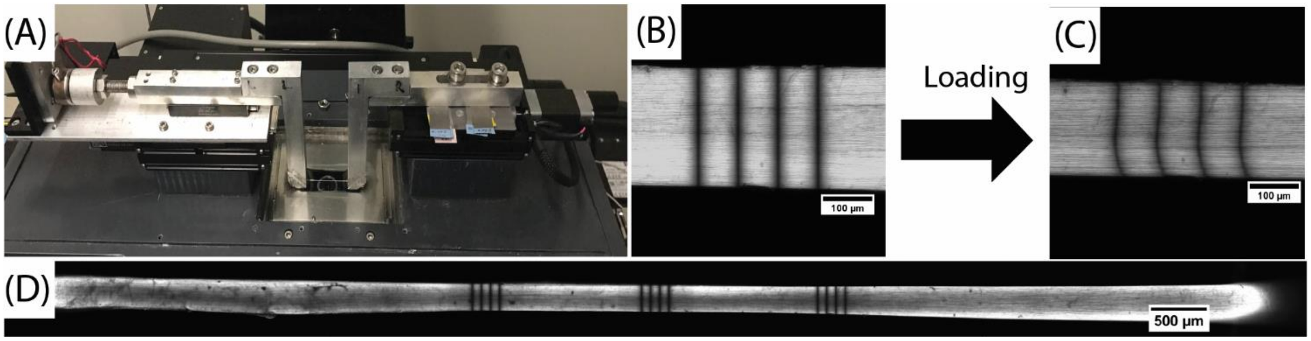


**Supplementary Figure 1.** Experimental setup for multiscale mechanical testing. (A) Custom uniaxial tensile testing device mounted atop a confocal microscope. (B & C) Representative images of a photobleached line (PBL) site at prior to and after loading, respectively. (D) Tendon sample held at a 10 mm gauge length with PBL sets at the sample center and ± 1.5 mm.

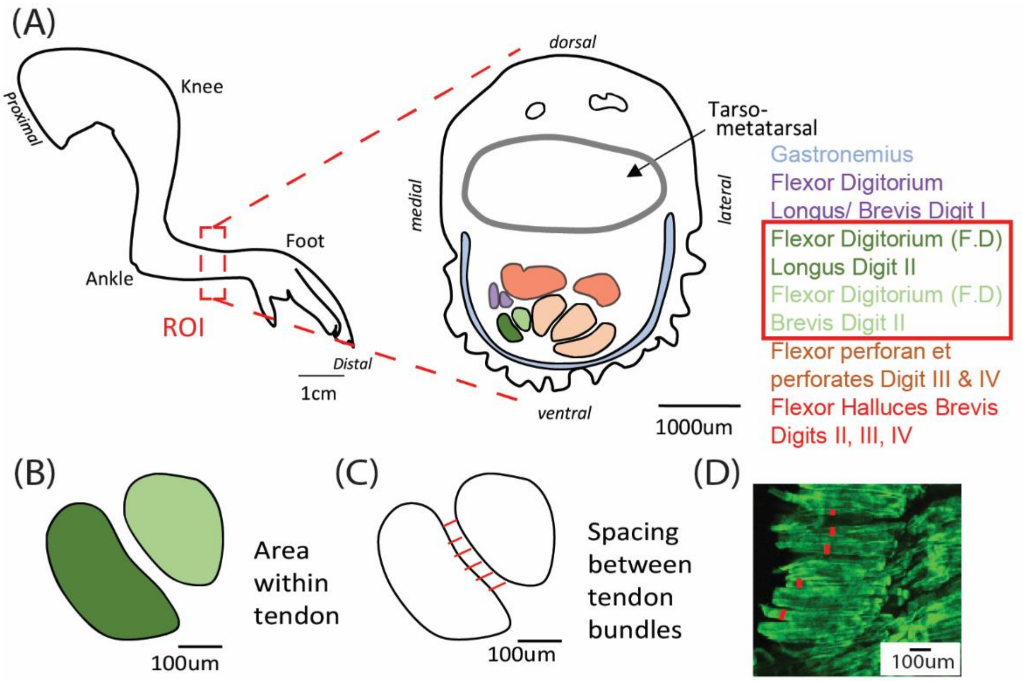


**Supplementary Figure 2.** (A) Schematic of a chick embryonic hindlimb showing a region of interest (ROI) through the tarsometatarsal region, and detailed morphology of cross-sections through this ROI with individual tendons labelled and color matched. (B) Schematic of the flexor digitorium longus and brevis digit II tendons to quantify the cross-sectional area within a tendon, (C) and the spacing between tendons, shown by red lines. (D) Confocal data of collagen binding protein (CNA35-eGFP) stained fibers were used to measure fiber diameter, indicated with individual red lines. All scale bars indicated.
